# Different energetic diets affect the maintenance of *Apis mellifera* L. colonies during off-season

**DOI:** 10.1101/629154

**Authors:** Gabriela Pinto de Oliveira, Samir Moura Kadri, Bruno Giovane Emilio Benaglia, Paulo Eduardo Martins Ribolla, Ricardo de Oliveira Orsi

## Abstract

The aim of this study wasto evaluate the best energetic foodforuse in the maintenance of honey bee colonies during the off-season. To do this, 20*Apis mellifera* beehives were used(with five beehives per treatment): CTL,control (without feeding); SJ,sugarcane juice; SS,sugar syrup; and IS,inverted sugar. We evaluated the food consumption, population development, and physiological state (expression of vitellogenin and hexamerin 70agenes)of eachcolony.The results showed that the supplementation of colonieswith sugar syrup resulted in an intermediateconsumption and thebetter development of the colony.In addition, this diet ensured that the colonies were in a good physiological state,as beesfed this diet presentedthe highest relative expression levels of vitellogenin and hexamerin 70ameasuredamong all thediets tested.Therefore, sugar syrup was concluded to be the best artificial energetic food for use in thesupplementation of honey bee colonies during the off-season.

## 1. Introduction

All of the nutritional needs of honey bees (*Apis mellifera*) for their complete development, maintenance, reproduction, and longevity need to be supplied through food sources available in the environment (Vaudo et al. 2015). In nature, *A. mellifera*bees meet all their nutritional needs through the ingestion of nectar and pollen (Degrandi-Hoffman and Chain 2015).Nectar has an undeniable importance to colonydevelopment, since it is the main source of energetic food forbees and permitstheir survivalthrough the off-season (Traynor et al. 2015). During the off-season, however, there is a shortage of blossomsand, consequently, of food for the bees. Thus, nutritional stress caused by food shortages or the availability of foods of only low nutritional value may lead to a reduction in the metabolic activity of the bees (Crailsheim 1990; Wang et al. 2016). It has also been observed that whenthere is little available natural food, there are reductions in the number of worker bees in the hive, queen’s layingand survival rates of individuals, and increases in escape or abandonment rates (Paiva et al. 2016).

This nutritional stress can cause alterations in thebees’ metabolic pathways that influence several biological processes, including the expressionof such genes as vitellogenin and hexamerin 70a. These genes are considered storage genes because they give rise to proteins that are produced during the larval stage and remain stored in the hemolymph and/or in the fat body (Martins et al. 2008).

Vitellogenin is important in the immune system and longevity of bees because this protein is a zinc carrier, and thus protects many cell lines against oxidative stress and apoptosis. Decreased expression of this gene product due to high expression levels of juvenile hormone was also previously shown to be related to reductions in the numbers of functional hemocytes in forager bees due to decreases in the quantity of zinc carried by vitellogenin(Falchuk 1998; Pinto et al. 2000;Amdam et al. 2004; Harwood et al. 2016).

Hexamerin 70a is a gene related to storage, which is expressed in the larval, pupal, and adult stages of workers, queens, and drones, and is the only protein with a hexamerin subunit present in significant quantities in the fat body and adult hemolymph of*A. mellifera*. The synthesis of hexamerins, specifically the subunit 70a, like thatof vitellogenin, is directly related to the quantity and quality of food intake. In older worker bees, such as forager bees, the levels of this protein are reduced due to the fact thatsuch bees have a low-protein diet, since they do not collect and consume much pollen and insteadpreferably consume nectar (Martins et al. 2008).

Beekeepersoften provide artificial food to their colonies in the off-season to lessen the negative effects that occur to bee colonies during this period in which food resources are drastically reduced, and thus ensure the survival and good performance of the colony.Therefore, the objective of this study was to select the best energetic food to be offered to bee colonies so that theirmaintenance is guaranteed during the off-season. This was done whiletaking into account the food consumption, population development, and physiological state of the colonyin terms of the expression of the genes for vitellogenin and hexamerin 70a when bee colonies were provided different foods.

## 2. Materials and methods

### 2.1 Field experiment

For the field experiment, 20 *Apis mellifera* beehives were selected, and the number of brooding and feeding frames in them were standardized. The food treatments used were the following: control (CTL), in which no artificial food was provided; feeding with sugarcane juice (SJ) produced by the research laboratory itself; feeding with sugar syrup (SS) prepared using pre-boiled filtered water and commercial crystal sugar; and feeding with inverted sugar (IS) purchased from Atrium Food Group, Campinas, Sao Paulo, Brazil.

Food was supplied twice a week in the amount of 1 L per hive for a period of 60 days by means of a Boardman artificial feeder. The experiment was carried out in June and July of 2015. During the experimental period, food consumption was measured weekly.

Population development, includingthe number and area of open and closed brooding areas in the central nest structure, was measured monthly in the hives throughout the experimental period usingthe methods used and described byLomele et al. (2010).

The number of brooding and feedingframes was quantified weekly by visual examination. Physicochemical analyses of the food provided were carried out at the Laboratory of Bromatology of the Experimental Farm Lageado, College of Veterinary Medicine and Animal Science, Universidade Estadual Paulista, UNESP, Botucatu, Sao Paulo, Brazil. Total sugar reduction was performed according to Welke et al. (2008), calorimetric and dry matter analysis according to Sodré et al. (2011), and ashing according to Sodré et al. (2007).

### 2.2 Genetic analysis

#### 2.2.1 Honeybee collection

The collection of bees for use in the analysis of the relative expression levels of the selected genes was done on day 0, which was used as experimental control, as well as on day 30, which was identified as “Moment 1” (M1) of the collection, and on day 60, which was designated as “Moment 2” (M2) of the collection. Five worker bees developing nursing behavior were collected from the central group, which were identified as nursing bees (N), and 5 worker bees were also collected that were carrying pollen in their corbicle, which in turn were identified as forager bees (F). During the experimental period, all ofthe colonies receiving the SJ treatment died, which made it impossible to collect these bees for analysis. After collection, the bees were immediately stored in a freezer at-80 °C for future RNA extraction.

#### 2.2.2 Relative expression of storage genes

To analyze the expression of vitellogenin and hexamerin 70a, 5 forager and 5 nursing bees were randomly collected and immediately frozen at −80 °C from each experimental colony at days 0, 30, and 60. Total RNA was extracted from the heads of 5 bees of each type (Scharlaken et al., 2008) using 500 μL of TRIzol® Reagent (Life Technologies) for each sample according to the manufacturer’s instructions. The extraction product was visualized on 1% agarose gel and quantified using aNanoDrop Instrument (ND-1000 Spectrophotometer). The RNA was treated with DNase I, incubated for 60 min at 37 °C and then for 10 min at 75 °C. A solution was then prepared of 0.75 mM oligo dT,0.15 mM random oligos,0.75 mM deoxynucleotide triphosphates, and 1 μLof RNA, and then it was incubated at 65 °C for 5 min, after which it was placed on ice for 1 min. We added 1× buffer dithiothreitol 0.005,RNaseOUT (40 U/μL), and 100 U SuperScript® III Reverse Transcriptase (Invitrogen) to this preparation. Complementary DNA synthesis was performed at 50 °C for 60 min, followed by 15 min at 70 °C.

Amplification was performed with real-time quantitative polymerase chain reaction (RT-qPCR) in a 25μL reaction mixture using the SYBR® Green PCR Master Mix (Applied Biosystems) and 0.2 μM of each primer. The sequences and details of the primers used are provided in Table I. The RT-qPCR reactions were performed using ABI 7300 (Applied Biosystems) equipment under the following conditions: 1 cycle at 50 °C for 2 min; 1 cycle at 94 °C for 10 min; and 40 cycles of 94 °C for 15 s and 60 °C for 1 min. The dissociation curve was obtained under the following conditions: at 95 °C for 15 s, 60 °C for 30 s, and 95 °C for 15 s. The determination of the expression levels of vitellogenin and hexamerin 70a was performed in triplicate, andthe expression of the actin gene was used as the control (Scharlaken et al. 2008). For each reaction, a negative control consisting of a mixture of reagents and water was also used.

**Table I:**
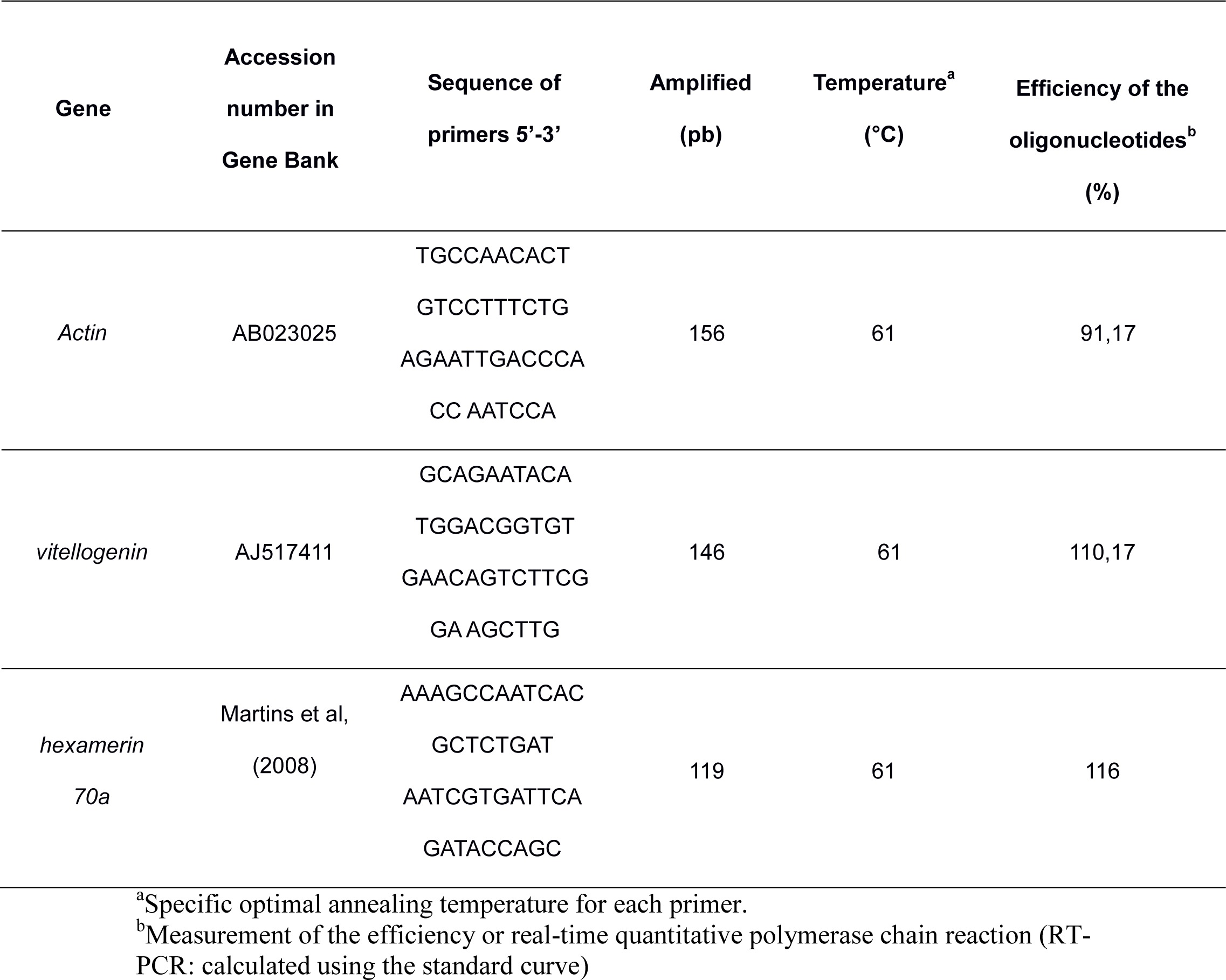
Oligonucleotides used in gene expression study of *Apis mellifera* that were fed with different energetic foods during the off-season.

The efficiency of the oligonucleotides (E) was calculated from 4 dilutions of complementary DNA samples (1:5, 1:25, 1:125, and 1:625) using the formula E = 10 (– 1/inclination). The quantification of a gene’s relative expression (R) was determined according to Pfaffl (2001), defining the crossing point as the point at which the detected fluorescence was appreciably above the background fluorescence and using the formula:

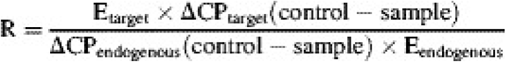

### 2.3 Statistical analyses

The data obtained for food consumption, population development, and gene expression were first tested for normality (Anderson-Darling test) and homogeneous variances (Levene’stest).When significant deviations (P<0.05) from these assumptions were detected, the data were compared using the non-parametric Mann-Whitney test, and the median and interquartile intervals (Q1_Q3) were presented. When no significant deviations from normality and homoscedasticity were detected, the data were analyzed with one-way ANOVA, and the mean ± standard deviation values were presented. P-values below 0.05 were considered significant. All statistical analyses were performed using Minitab statistical software.

## 3. Results

On mean ± standard deviation, 0.994.8 ± 310, 894.6 ± 291, and 433.9 ± 227.6 mL of SJ, SS, and IS, respectively, were consumed weekly. The CTL, SJ, SS, and IS treatments underwent 100, 75, 0, and 0% colony losses, respectively.

The results of the physicochemical analyses of the different foods are presented in Table II. Significant differences were observed in the analysis ofSJ ash (0.27 ± 0.02%),in that this value differed significantly from that for SS (0.01 ± 0.001%), but not from that for IS (0.11 ± 0.04%).

**Table II.** Physicochemical analyses of different energetic foods (sugarcane juice, sugar syrup and inverted sugar).

|  | Ashing (%) | Calorimetric<br>(Kcal Kg<sup>-1</sup>) | Drymatter (%) | Reduced sugars<br>(%) |
| --- | --- | --- | --- | --- |
| SugarcaneJuice | 0.27±0.02a | 3903.0 | 15.94±0,00a | 21.15±1,6a |
| Sugar syrup | 0.01±0.00b | 4155.0 | 53.84±0.41b | 41.52±2.8b |
| Inverted sugar | 0.11±0.04ab | 3895.0 | 75.66±0.75c | 0.82±0.0c |

Different lowercase letters on the same line indicate statistical differences between means (p <0.05).

The calorimetric analysis of the foods showed that SS presented the highest energetic value (4155.0 kcal kg^-1^) of all the foods tested (3903.0 kcal kg^-1^forSS and 3895.0 kcal kg^-1^forIS).The dry matter values ofSJ and SS were lower than that of IS, indicating they had higher moisture content. The analysis of the total reducing sugars in each food type showed a higher value for SS (41.52 ± 2.8%), which differed significantly from those found for SJ (21.15 ± 1.6%) and IS (0.82 ± 0.01%).

The data contained in Table III represent the number of brooding and feeding frames in colonies under different treatments. The number of brooding frames was higher for colonies in theSS and IS treatments compared to that in the CTL and SJ treatments. However, the number of feedingframes showed no differences among treatments.The data shown in Table IV represent the areas of open and closed brooding areas (cm^2^) observed in colonies subjected to different treatments. The treatments SS and IS presented larger closed brooding areas than the other treatments did. For the open brooding areas, the largest observed area was recorded in theIS treatment.

**Table III.**
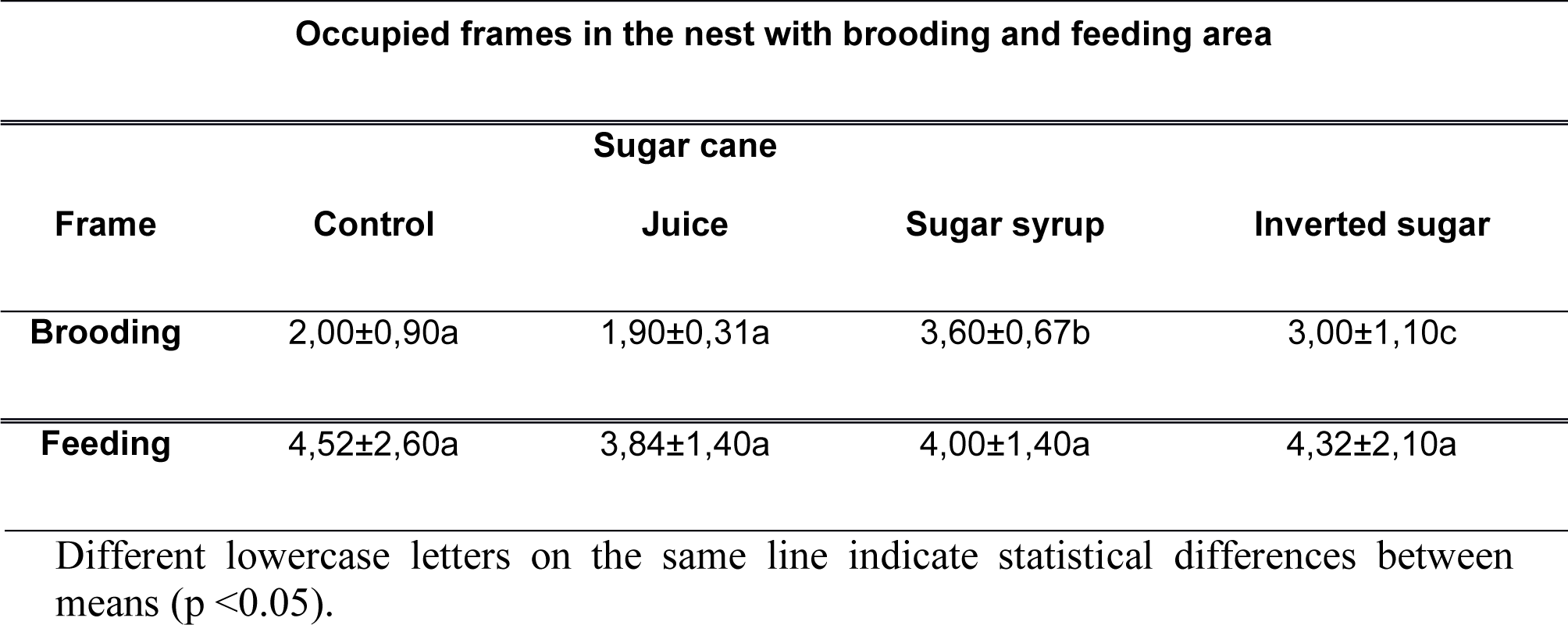
Mean and standard deviation of the number of tables occupiedi n the nest containing brooding and feeding frames for control, sugarcane juice, sugar syrup and inverted sugar treatments during the experimental period.

**Table VI.**
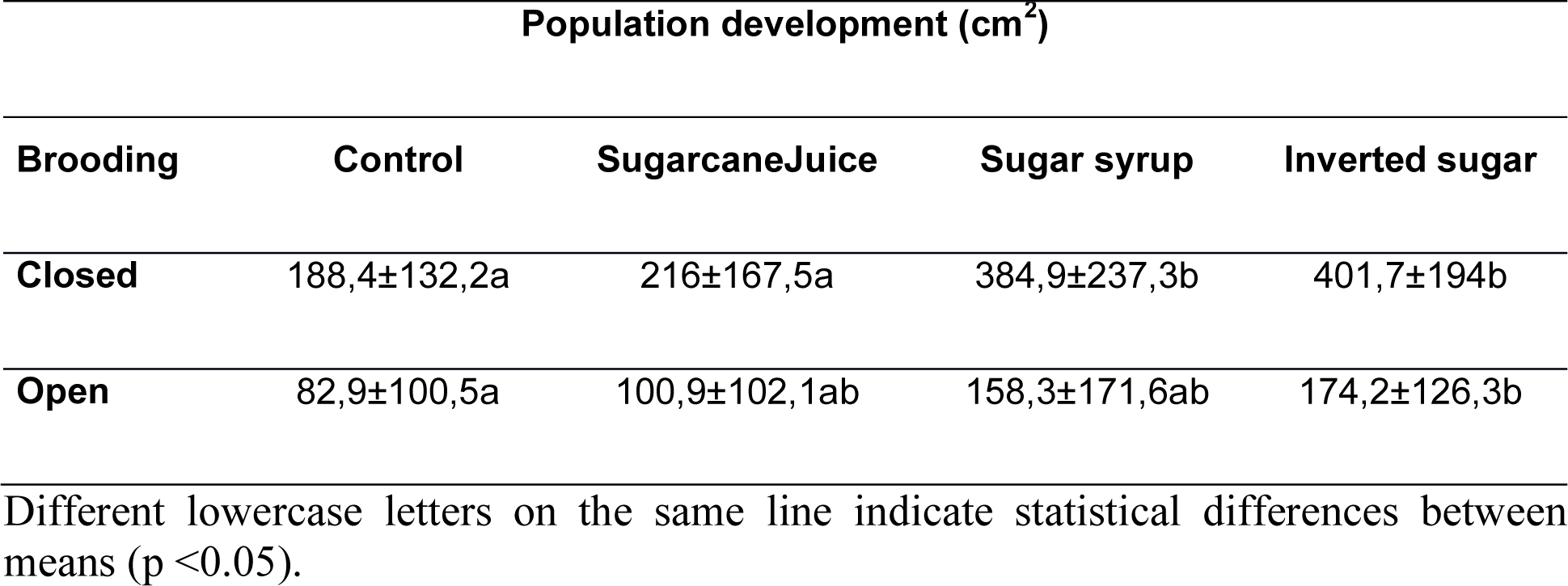
Mean and standard deviation of the open and closed (cm^2^) brooding area referring to the treatments control, sugarcane juice, sugar syrup and inverted sugar, during the experimental period.

The mean and standard deviation of the relative expression levels of the vitellogenin and hexamerin 70a genes in each treatment were calculated, and these are presented in TablesI and II and Figures 1 and 2.

**fig. 1.**
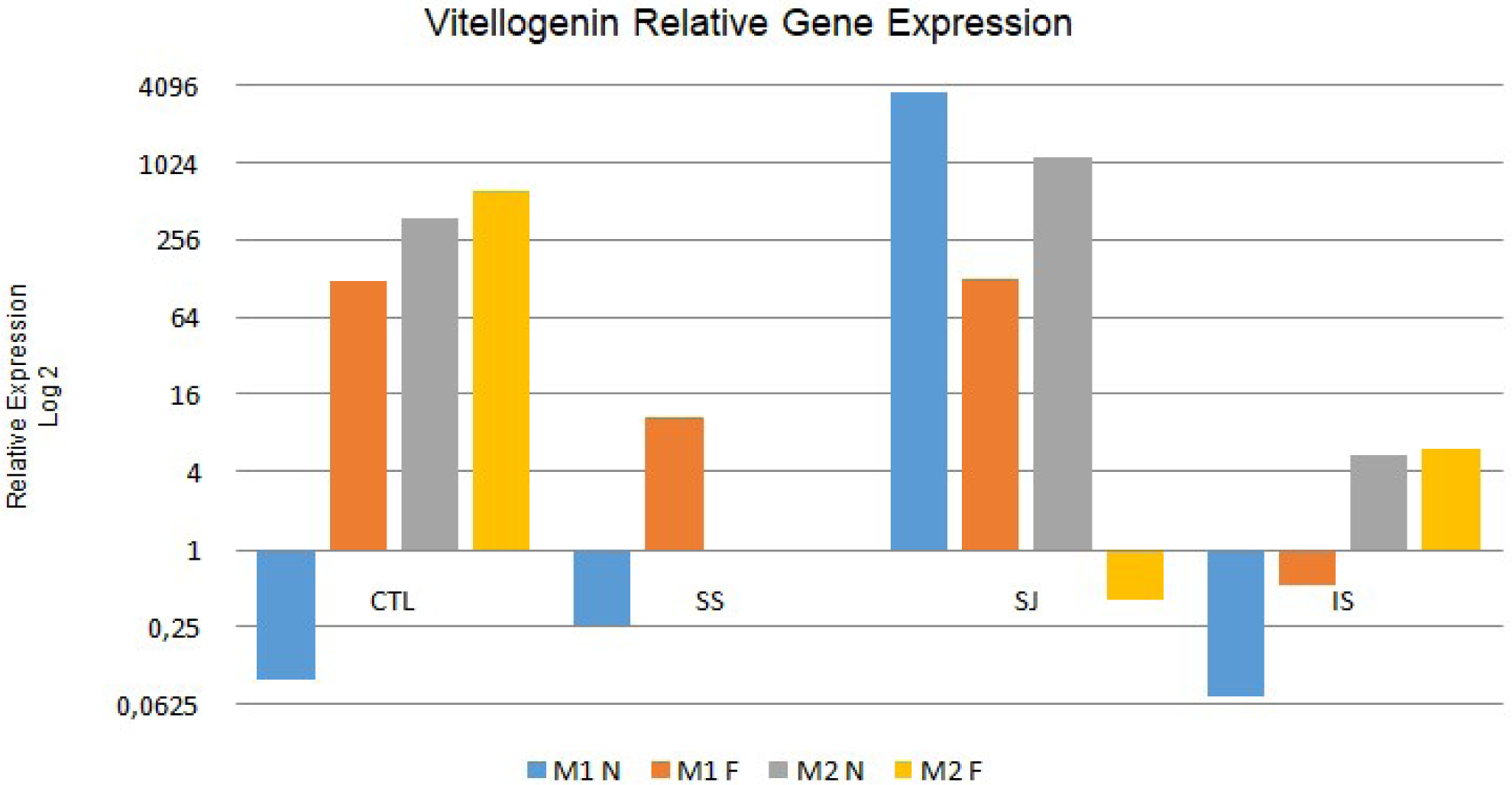
Relative expression of the vitellogenin gene in nursing bees (N) and forager bees (F) of the different treatments used after 30 days (Moment 1 -M1) and 60 days (Moment 2 -M2). CTL: control; SJ: sugarcane juice; SS: sugar syrup; IS: Inverted sugar. M2 N and M2 F: Data not obtained due to death of the colonies during the experimental period

**fig. 2.**
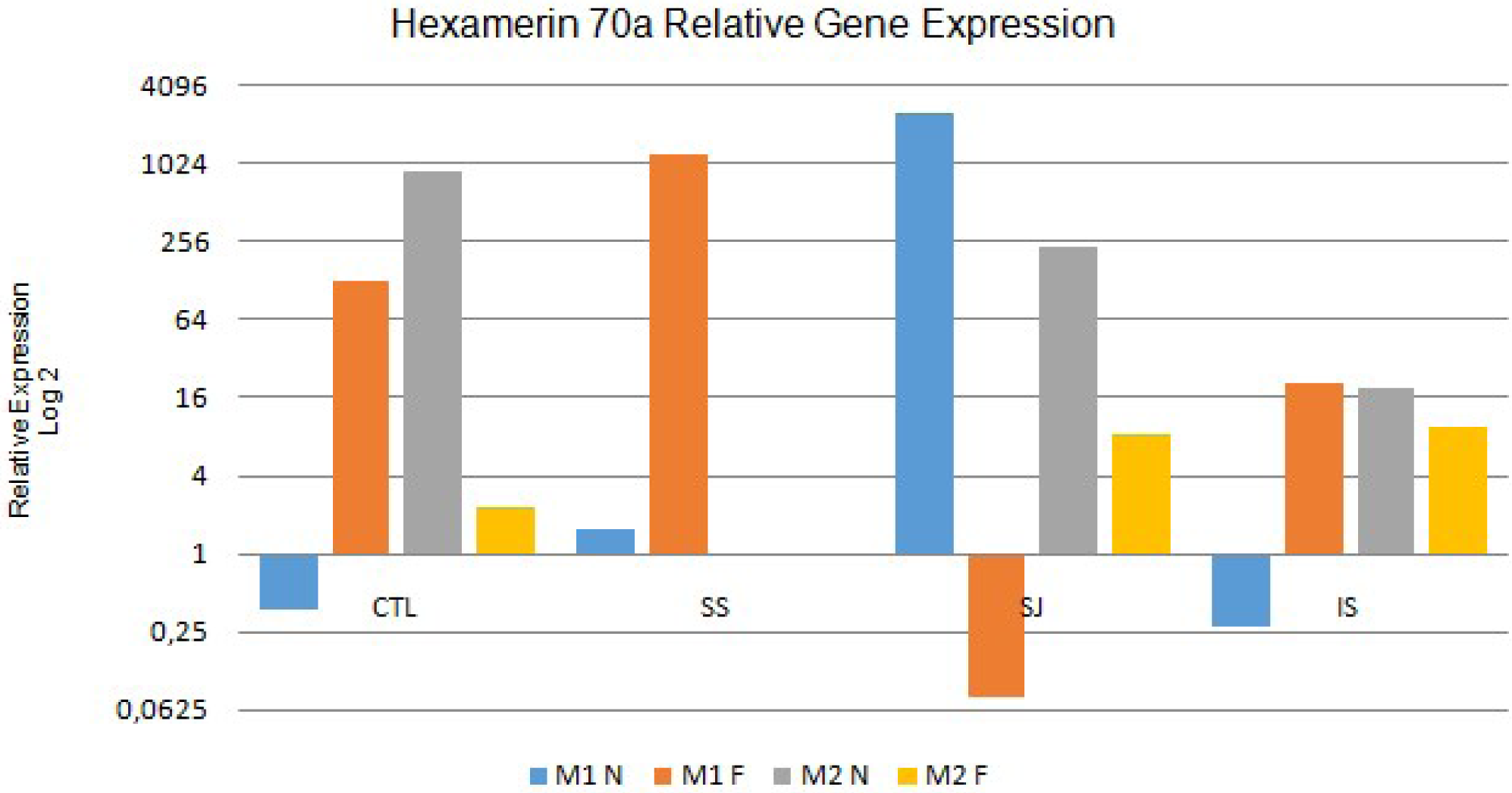
Relative expression of the hexamerin 70a gene in nursing bees (N) and forager bees (F) of the different treatments used after 30 days (Moment 1 -M1) and 60 days (Moment 2 -M2). CTL: control; SJ: sugarcane juice; SS: sugar syrup; IS: Inverted sugar. M2 N and M2 F: Data not obtained due to death of the colonies during the experimental period

In terms of the relative expression of the vitellogenin gene, from the comparison between nursing and foragerbees at M1 and M2 (Table V and Figure 1) it can be observed that the nursing bees in the CTL, SJ, and IS treatments at M1 and in the CTL and IStreatments at M2 presented significantly lower relative expression levels of this gene (P<0.05). However, the inverse was observed in the bees analyzed from the SS treatment at both M1 and M2 (P <0.05), with the nursing bees expressingthis gene at levels 2862 times greater than those in the foragerbeesat M2.

**Table V.**
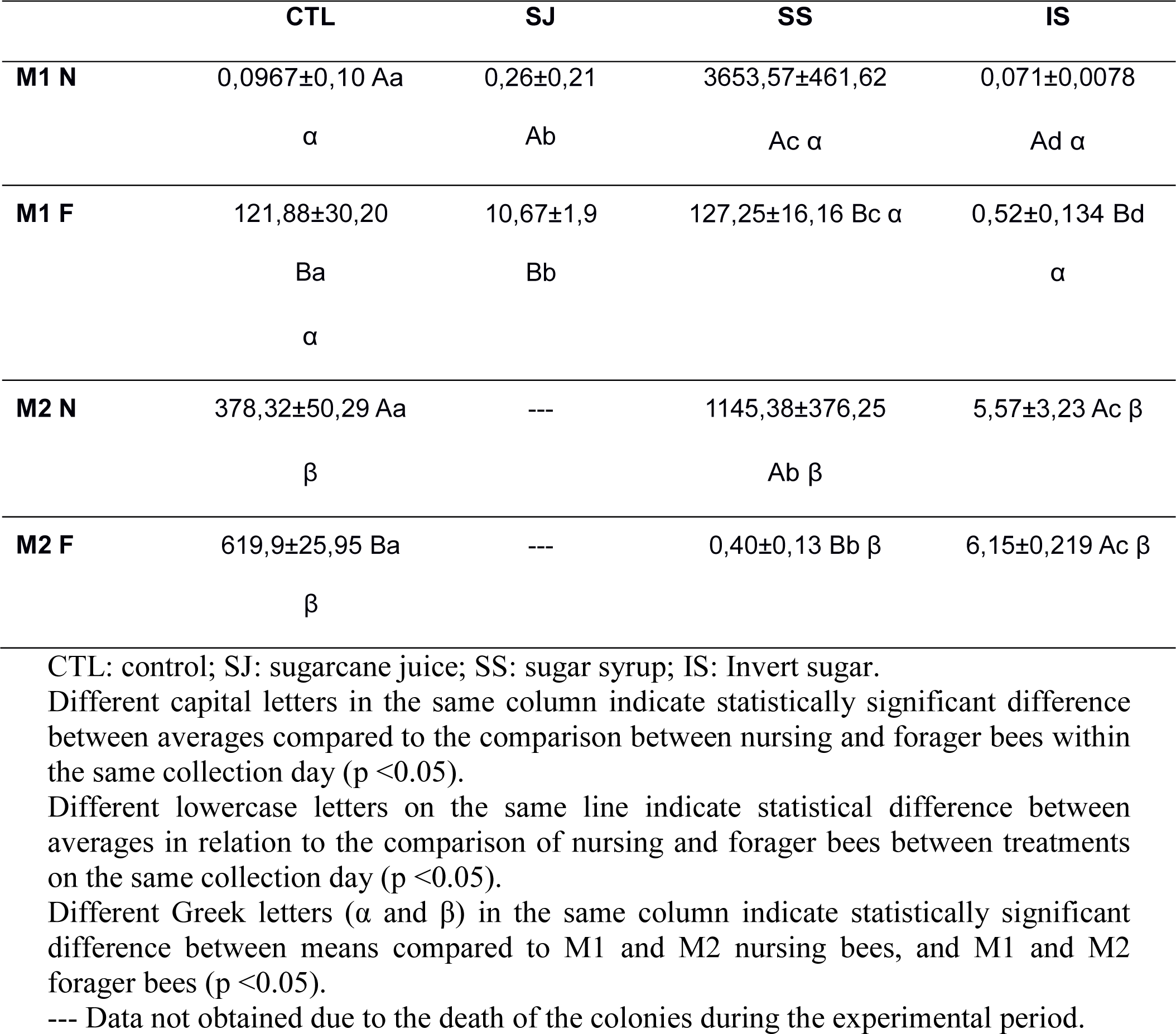
Relative expression of the vitellogenin gene in nursing bees (N) and forager bees (F) of the different treatments used after 30 days (Moment 1 -M1) and 60 days (Moment 2 -M2).

At M1, when analyzing only the nursing bees in relation to the diets provided, the treatments CTL, SJ, and IS resulted in the downregulation of this gene (i.e. a decrease in the relative expression level of the gene in comparison to that in the control group), whereas in the SStreatment the upregulation of this gene (i.e. increased relative expression) was noted. For the foragerbees sampled at the same time, only those treated with IS presented downregulation of this gene, which showedthat there were differences in the patterns of gene expression between the nursing and forager bees.

When analyzing nursing bees at M2, the expression of this gene was upregulated in bees in all treatments, although the CTL and SS treatments presented relatively higher levels of expression compared to those in the IStreatment (67,92 and 205,63 times more than in the IS treatment, respectively). It was observed that bees fed with SS showed greaterexpression of this gene in comparison to those in the other treatments, similarly to what was noted above for the nursing bees collected at M1. However, for the foragerbees collected at the same time, only those treated with SS presented downregulation of this gene in relation to the control.

When changing the focus of the data analysis and comparing the results obtained for nursing andforager bees between M1 and M2 in the different treatmentsto check for changes in the expression pattern of this gene after 60 days of feeding, it was observed that all the analyzed treatments showed differences in their relative expression levels (P<0.05). In the CTL and IS treatments, both nursing andforager bees showed increasedexpression of the vitellogenin gene during the experiment (P<0.05). However, with the SS treatment, the inverse occurred and there was greater relative expression at M1 than at M2 in both nursing and forager bees.

For the comparison between nursing and forager bees at M1 and M2 in relation to their relative expression levels of the hexamerin 70a gene (Table VI and Figure 2), in all treatments except SS there were lower relative expression levels of this gene in the nursing bees than the in the foragerbees at M1 thismeantthat in the nursing beesthis gene was downregulated, whereas in the forager bees it was upregulated. However, the inverse patternoccurred in the SStreatment, in which the nursing bees presented greater relative expression levels of hexamerin 70a than the forager bees did, and in the nursing bees a very extensive upregulation was observed; specifically, the expression of this gene reached a value 33483 times higher than that in nursing bees at M1, whereas in the forager bees this gene’s expression was downregulated.

**Table VI.** Relative expression of the hexamerin 70a gene in nursingbees (N) and forager bees (F) of the different treatments used after 30 days (Moment 1 -M1) and 60 days (Moment 2 -M2).

|  | CTL | SJ | SS | IS |
| --- | --- | --- | --- | --- |
| **M1 N** | 0,3818±0,02 Aa<br>$\alpha$ | 1,557±0,34 Ab | 2554,82±243,59<br>Ac $\alpha$ | 0,2689±0,02<br>Ad $\alpha$ |
| **M1 F** | 125,46±2,64 Ba<br>$\alpha$ | 1217,16±383,91 Bb | 0,0763±0,003 Bc $\alpha$ | 20,818±1,43 Bd<br>$\alpha$ |
| **M2 N** | 911,30±49,52 Aa<br>$\beta$ | --- | 233,83±30,94<br>Ab $\beta$ | 19,256±1,78 Ac<br>$\beta$ |
| **M2 F** | 2,276±0,21 Ba $\beta$ | --- | 8,437±1,04 Bb $\beta$ | 9,531±0,60 Bb $\beta$ |

CTL: control; SJ: sugarcane juice; SS: sugar syrup; IS: Invert sugar. Different capital letters in the same column indicate statistically significant difference between averages compared to the comparison between nursing and forager bees within the same collection day ( $p < 0.05$ ). Different lowercase letters on the same line indicate statistical difference between averages in relation to the comparison of nursing and forager bees between treatments on the same collection day ( $p < 0.05$ ). Different Greek letters ( $\alpha$ and $\beta$ ) in the same column indicate statistically significant difference between means compared to M1 and M2 nursing bees, and M1 and M2 forager bees ( $p < 0.05$ ). --- Data not obtained due to the death of the colonies during the experimental period.

At M2, this gene was upregulated in both the foragerand nursing bees in all treatments. At this time, it was also possible to observe thatthe nursing bees had relatively higher levelsof expression of the hexamerin 70a gene than the foragerbees didin all treatments.

## 4. Discussion

Feeding honey bee colonies with artificial energetic foods during the off-season ensures the correct annual operation of the colony. For this to be effective, it is necessary to choose the best energetic food to offerthe bees to guaranteetheproper development of a colony for the producer.The lower consumption of inverted sugar in the present study may suggest that this energy supply was less attractive to the bees when compared to that offered in the other treatments. This fact can be attributed to the higher viscosity of the inverted sugar, which was supplied to the hives at its full concentration (Gratão et al. 2004).

The dry matter data followed opposing trends among treatments to thedifferent food types’ viscosities, since the moisture content of the food is inversely proportional to its viscosity (Yanniotis et al. 2006; Cui et al. 2008). Therefore, sugar syrup and sugarcane juice had lower viscosities, which may have favored their consumption since nectar, a natural energetic food of bees, has a low viscosity and high humidity (Garcia et al. 2005; Hazlehurst and Karubian 2016). Furthermore, the higher reducing sugarand caloriccontent and the lower ash content of sugar syrup detected in the analyses carried out in this study, along with this food having an adequate dry matter content, indicated that this was the food that we supplied to the bees that most closely resembled honey, which is the main natural source of energy reserves for bee colonies. Thus, because it has a composition closer to that of the bees’ natural food, it was, at least in bromatological terms, the best source of artificial food for bees that was tested in this study.

Castagnino et al. (2006) showed that energetic supplementation during the off-season increases the queen’s posture. In addition to supplying an energetic diet, a protein diet is also essential for colonymaintenance and improving the queen’s posture (Frias et al. 2016).However, under the conditions of this experiment, the colonies had bee bread reserves,and thus no protein supplementation was required. In this case, the energetic supplementation provided assisted in the maintenance of the colonies, and was able to explainthe greater number of brooding frames observed in the SS and IS treatments, suggesting that these energetic foods provided the necessary energetic support for the queen’s posture during this period. Castagnino et al. (2006) obtained a large brooding area incolonies fed with sugar syrup, which was similar to the results of the present study. The energy provided by the consumption of the sugar syrup and inverted sugar probably stimulated the queen’s laying.

The loss of all of the colonies subjected to the SJ treatment over the experimental period probably occurred because the sugarcane juice (the diet offered to bees in the SJ treatment) may have undergone fermentation at ambient temperature (Pedroso et al. 2005).Given this, it was not possible to obtain data on the relative expression of the tested genes at Moment 2 in the nursing and forager bees in this treatment.

The analysis of nursing bees at M1 demonstrated an upregulation in vitellogenin expression in the SS treatment only. This shows that sugar syrup had a more direct influence on nursing bees than the other foods provided throughout the experimental period. This result may be related to several factors, such as food viscosity, energetic value, and the maintenance of foodintegrity and quality at room temperature.

The vitellogenin expression levels observed after 60 days of feedingsuggested that the SS treatment had a greater influence on the expression of this gene than the other diets, and it also facilitated better population development of the colony since the values of almost all of the performance parameters observed were higher in this treatment compared to those in the other treatments. The forager bees, which live forapproximately 21 days, presented less relative expression of this gene than the nursing bees, which were less than 15 days old, at both M1 and M2. This possibly occurred due to the fact that there was a higher concentration of juvenile hormone in the hemolymph of the older bees, which may have influenced their biosynthesis of vitellogenin. As noted earlier, vitellogenin is a protein that is related to the prevention of oxidative stress since it is a zinc carrier, andthe occurrence of low levels of this protein can compromise the immune system (Dallacqua et al. 2007).

Therefore, the results of this study demonstrated that the supplementation of honey bee colonies in the field during the off-season with sugar syrup resultsin an intermediate level of consumption of this food by them and greater colony development, and also ensured that the bees were in a better physiological state. This showed that this wasthe most beneficial artificial energetic food tested in this study.

## 5. Acknowledgements

Fundação de Amparo à Pesquisa do Estado de São Paulo (FAPESP) for the scientific initiation scholarship provided (FAPESP process: 2017/12001-2).

